# The Il1r2-CreERT2 knock-in mouse enables inducible labeling of slow-cycling epidermal basal cells and genetic ablation of the IL-1 decoy receptor

**DOI:** 10.64898/2026.05.25.727717

**Authors:** Hung Manh Phung, Nguyen Thi Kim Nguyen, Ikuto Nishikawa, Naoki Takeda, Takeru Fujii, Guangqi Gao, Yasuyuki Ohkawa, Kimi Araki, Aiko Sada

## Abstract

The interfollicular epidermis is maintained by spatially organized basal cell populations with distinct molecular signatures and division kinetics; however, the markers that define these populations remain poorly defined. In this study, we identified *Il1r2*, which encodes the IL-1 decoy receptor IL-1R2, as a marker of the slow-cycling basal population in the epidermis. Single-cell RNA-seq analysis of epidermal and hair follicle basal populations in the murine tail skin revealed that *Il1r2* is preferentially expressed in the slow-cycling epidermal basal population, and immunofluorescence staining confirmed its protein localization in tissues. To enable fate mapping of this population, we generated an Il1r2-CreERT2 knock-in mouse line using a CRISPR-Cas9-based PITCh method. Tamoxifen induction in Il1r2-CreERT2/Rosa-tdTomato mice exhibited selective labeling of basal cells localized to the slow-cycling interscale region of the tail epidermis. Because the CreERT2 cassette was inserted into the *Il1r2* coding sequence, homozygous Il1r2-CreERT2 knock-in mice can also serve as an *Il1r2* knockout model through targeted gene ablation. Thus, the Il1r2-CreERT2 mouse line provides a dual genetic tool for lineage tracing of the slow-cycling epidermal basal population and for functional modulation of IL-1 signaling *in vivo*.

## Introduction

The interfollicular epidermis is maintained by the self-renewal and differentiation of epidermal stem cells residing in the basal layer. Recent advances in single-cell RNA sequencing (scRNA-seq), lineage tracing, and mathematical modeling have revealed that basal cells are not homogeneous but comprise subpopulations with distinct molecular signatures and cellular behaviors (Ghuwalewala et al., 2024; Sada et al., 2016; Vu et al., 2022). Similar basal cell heterogeneity has been reported in the human epidermis, where spatial organization along interridge and rete ridge structures is associated with distinct basal cell states (Cheng et al., 2018; Ghuwalewala et al., 2022; Ishikawa et al., 2026; Negri & Watt, 2022; Wang et al., 2020). However, despite these advances, molecular markers that can robustly define distinct epidermal basal cell populations remain limited.

The mouse tail epidermis serves as a model for studying basal cell heterogeneity. This tissue is organized into spatially distinct epidermal regions: the interscale and the scale. Hair follicles and pigmented scale regions are arranged in a regular, repetitive pattern along the tail, forming a characteristic scale-like structure (Gomez et al., 2013)(Fig. 1A). The interscale region occupies the areas between scales where hair follicles penetrate the epidermis. These regions can be distinguished by their differentiation lineages: interscale regions undergo orthokeratotic differentiation (K10^+^ area), whereas scale regions exhibit parakeratotic differentiation (K31^+^ or K36^+^ area), allowing spatial visualization by whole-mount immunostaining.

**Figure 1.**
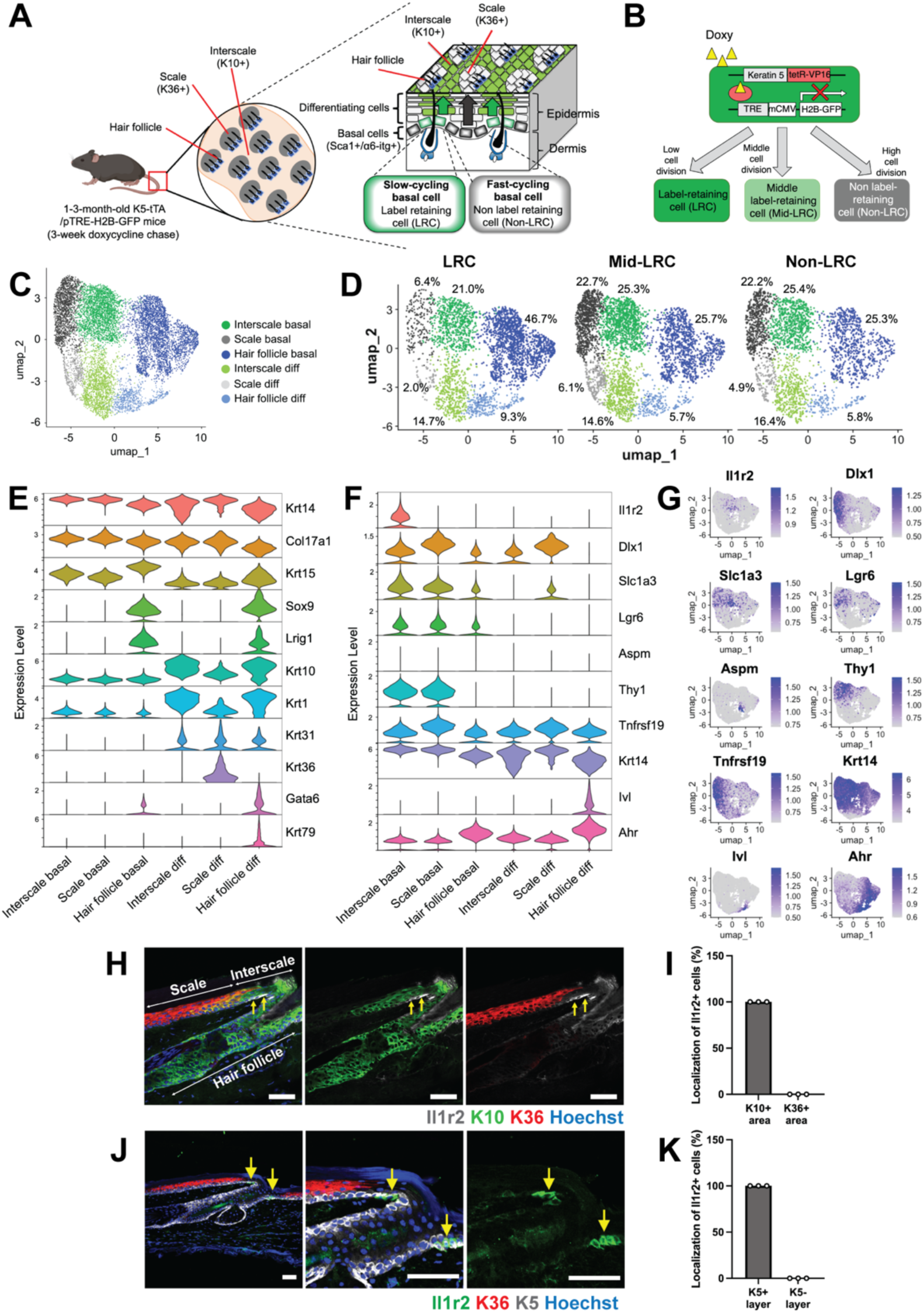
scRNA-seq analysis reveals *Il1r2* expression in a slow-cycling epidermal basal population. **(A)** Schematic representation of mouse tail skin. Slow-cycling epidermal basal cells give rise to the K10+ interscale lineage (green), whereas fast-cycling epidermal basal cells give rise to the K36+ scale lineage (gray). **(B)** Schematic of the K5-tTA/pTRE-H2B-GFP mouse model used for label-retaining analysis. **(C, D)** UMAP plots showing cell clusters identified based on established marker gene expression and the distribution of LRC, mid-LRC, and non-LRC cells within each cluster. Interscale diff: interscale differentiating cells; scale diff: scale differentiating cells; hair follicle diff: hair follicle differentiating cells. **(E, F)** Violin plots showing ALRA-imputed expression of marker genes used to define basal and differentiating cell clusters (E) and epidermal stem/progenitor markers (F). **(G)** Feature plots showing ALRA-imputed expression of representative epidermal stem/progenitor markers across clusters. **(H)** Immunostaining of tail skin sections for Il1r2, K10 (interscale marker), and K36 (scale marker). **(I)** Quantification of the percentage of Il1r2^+^ cells localized within the K10^+^ interscale and K36^+^ scale regions. **(J)** Immunostaining of tail skin sections for Il1r2, K36 (scale marker), and K5 (basal layer marker). **(K)** Quantification of the percentage of Il1r2^+^ cells localized within the K5^+^ basal and K5^−^ suprabasal layers. Yellow arrows indicate Il1r2^+^ cells. Scale bars: 50 µm. Each dot represents one mouse. Il1r2^+^ cells are exclusively and consistently localized to the interscale basal layer across all samples.

At the cellular level, these regions also differ in their proliferative dynamics. Studies employing the histone H2B-GFP label-retention assay have shown that the interscale region is enriched with infrequently-dividing, label-retaining cells (LRCs), whereas the scale region contains more frequently-dividing, non-LRCs (Sada et al., 2016)(Fig. 1A). Single-cell RNA-seq analyses using LRC and non-LRC basal populations have identified candidate markers, including Sox6, Vamp1, and Col17a1 for the LRC region, and Slc1a3, Krt15, and Sostdc1 for the non-LRC region (Ghuwalewala et al., 2022); however, these genes often exhibit variable or overlapping expression across multiple basal cell populations.

Several CreER-based lineage tracing tools have been developed to label distinct epidermal basal populations (Table 1). However, most of these lines lack the specificity required to clearly distinguish spatially defined basal cell populations. For example, although Ah-, K14-, and Inv-CreER were originally used for clonal analyses, they show global labeling without regional biases (Clayton et al., 2007; Gomez et al., 2013; Mascré et al., 2012). Dlx1- and Slc1a3-CreER preferentially label LRC and non-LRC populations in the interscale and scale regions, respectively; however, both lines exhibit cross-labeling between these regions (Sada et al., 2016). Similarly, the Lgr6- and Troy-CreER lines target the interscale epidermis, but also label additional structures in the hair follicles (Kretzschmar et al., 2016, 2021). Other lines, such as CD4-, Aspm-CreER, mark progenitors across both interscale and scale regions (Brandes et al., 2025; Ghuwalewala et al., 2024). Although Thy1 protein is enriched in the interscale, lineage-tracing data using Thy1-CreER have not been reported in the tail skin (Koren et al., 2022). Thus, there is no CreER driver with sufficient regional specificity to selectively and reliably label interscale (slow-cycling or LRC) versus scale (fast-cycling or non-LRC) basal cell populations.

Previous studies have suggested *Il1r2* (Interleukin-1 receptor type 2) as a candidate marker enriched in the epidermal LRC population (Ishikawa et al., 2026; Sada et al., 2016). IL-1R2 functions as a decoy receptor that sequesters IL-1α and IL-1β, thereby limiting the excessive activation of IL-1 signaling (Luís et al., 2022; Peters et al., 2013; Rauschmayr et al., 1997); however, its roles in skin homeostasis and epidermal stem cell regulation have not been reported.

In this study, we show that *Il1r2* is preferentially expressed in interscale basal cells, as revealed by scRNA-seq and antibody staining. To enable selective labeling and functional analysis of this population, we generated CreERT2 knock-in mice using a CRISPR-Cas9–mediated Precise Integration into Target Chromosome (CRIS-PITCh) method (Nakade et al., 2014; Sakuma et al., 2016). Because the CreERT2 cassette is inserted into the endogenous *Il1r2* locus, homozygous mice also function as *Il1r2* knockout mice. Thus, this model represents a genetic tool for lineage tracing slow-cycling epidermal basal cells and the genetic ablation of *Il1r2*.

## Material and Methods

### Analysis of scRNA-seq data

Publicly available scRNA-seq data (GEO accession: GSE205746) (Ghuwalewala et al., 2022) were reanalyzed. In the original study, LRC, mid-LRC, and non-LRC populations were isolated from α6-integrin⁺/Sca1⁺ epidermal basal fraction of tail skin following a 3-week doxycycline chase in K5-tTA/pTRE-H2B-GFP mice at 1–3 months of age.

Raw count matrices generated by the 10× Genomics platform were analyzed in R using Seurat (Satija et al., 2015). Cells with 200–5,000 genes and <10% mitochondrial gene content were used for downstream analyses (Fig. S1A). LRC, mid-LRC, and non-LRC datasets were normalized, scaled, and subjected to principal component analysis (PCA), followed by Harmony-based batch correction across biological replicates (Butler et al., 2018; Korsunsky et al., 2019). Graph-based clustering (resolution = 0.3) and Uniform Manifold Approximation and Projection (UMAP) (McInnes et al., 2018) visualization were then performed using dimensions 1–10 of the Harmony-corrected embeddings. A cluster with low median nFeature_RNA and nCount_RNA values relative to the remaining clusters was excluded. The filtered dataset was reprocessed using the same workflow (dims = 1:10, resolution = 0.3), yielding eight transcriptionally distinct clusters (Fig. S1B). Initial cluster identities were assigned based on the normalized expression of canonical marker genes (Fig. S1C, D).

Clusters corresponding to proliferating cells and infundibulum cells were excluded from subsequent analyses. The remaining cells were reprocessed using the same workflow with dimensions 1–10 of the Harmony-corrected embeddings and a higher clustering resolution (resolution = 0.5). Adaptively thresholded low-rank approximation (ALRA) imputation was performed using SeuratWrappers v0.4 (https://github.com/satijalab/seurat-wrappers) to impute dropout values and aid in the visualization and evaluation of marker gene expression patterns (Linderman et al., 2022). Final cluster annotation was performed based on ALRA-imputed expression patterns of established epidermal and hair follicle lineage markers, together with the relative abundance of LRC, mid-LRC, and non-LRC cells within each cluster, yielding 6 clusters (Fig. 1): interscale basal cells, scale basal cells, hair follicle basal cells, interscale differentiating cells, scale differentiating cells, and hair follicle differentiating cells. All analyses were performed in R using Seurat (v5.4.0) and Harmony (v1.2.4).

### Mice

C57BL/6J wild-type mice were purchased from Charles River Laboratories or Japan SLC, Inc. Mice of both sexes were used. The animals were housed in specific-pathogen-free conditions with a 12-hour light/dark cycle and *ad libitum* access to food and water. All animal experiments were approved by the Animal Care and Use Committee of Kumamoto University and the Animal Experiment Committee of Kyushu University, and the experiments were conducted in accordance with institutional guidelines.

### Generation of Il1r2-CreERT2 knock-in mice

An Il1r2-CreERT2 knock-in mouse line was generated using CRIS-PITCh technology (Sakuma et al., 2016) via the zygote microinjection of CRISPR-Cas9 components and a donor construct, as previously described (Tanimoto et al., 2022). A CreERT2 cassette followed by a β-globin polyadenylation (polyA) signal was inserted at the translational start site of exon 1 of the *Il1r2* locus, thereby disrupting the endogenous coding sequence.

For the donor construct, a ∼2.2kb CreERT2-polyA cassette was amplified from the pCAG-CreERT2 vector (Addgene #14797)(Matsuda & Cepko, 2007) and flanked by Il1r2 microhomology arms. Guide RNAs targeting the *Il1r2* locus and a generic PITCh-gRNA were designed using the PITCh Designer (Nakamae et al., 2017). All primer and gRNA sequences are listed in Tables 2 and 3.

Genome editing was performed in C57BL/6J zygotes by the Center for Animal Resources and Development (CARD) at Kumamoto University. The gene-editing mixture containing Cas9 nuclease (IDT), an *Il1r2* gene-specific gRNA (IDT), a PITCh-gRNA (IDT), and the donor construct was microinjected into fertilized eggs. Injected embryos were transferred into pseudopregnant females. Founder mice were screened by allele-specific PCR genotyping using primers flanking the insertion site, and correct targeting was further confirmed by genome sequencing across both 5′ and 3′ junction regions. The mice were backcrossed and maintained on a C57BL/6J background. Heterozygous mice were intercrossed to obtain homozygous Il1r2-CreERT2 mice. For lineage tracing, mice carrying Il1r2-CreER were crossed with Rosa-tdTomato reporter mice (Jackson Laboratory, Stock no. 007905).

### Tamoxifen injection

Tamoxifen (Sigma) was dissolved in corn oil at 10 mg/mL by heating. Il1r2-CreERT2/Rosa-tdTomato mice at 2–3 months of age were injected intraperitoneally at a single dose of tamoxifen (100 μg/g body weight). Skin tissue was collected at 2 weeks post-induction.

### Hematoxylin and eosin (H&E) staining

Tail skin samples were embedded in OCT compound (Sakura Finetech) and cryosectioned at 10 μm. Following fixation in 4% paraformaldehyde (PFA) for 10 minutes at room temperature (RT), the sections were stained with hematoxylin for 3 minutes and eosin Y for 15 seconds. Subsequently, the sections were dehydrated and mounted using Entellan (Merck Millipore). Images were acquired using an EVOS M5000 Imaging System (Thermo Fisher Scientific).

### Immunostaining of tail skin sections

Tail skin samples were embedded in OCT compound and cryosectioned at 10 µm. The sections were fixed in 4% PFA for 10 minutes at RT, and blocked for 1 hour in PBS containing 1% bovine serum albumin (BSA), 2.5% normal donkey serum (NDS), 2.5% normal goat serum (NGS), 2% gelatin, and 0.1% Triton. Subsequently, the sections were incubated overnight at 4°C with primary antibodies against K10 (Abcam, 1:500), K36 (Proteintech, 1:300), K5 (BioLegend, 1:1000), and Il1r2 (BD Biosciences, 1:100), followed by Alexa Fluor-conjugated secondary antibodies (Invitrogen, 1:300) for 1 hour at RT. Nuclei were counterstained with Hoechst 33342 (Sigma-Aldrich, 1:1000). Images were acquired using Nikon A1 and Zeiss LSM 900 confocal microscopes.

### Whole-mount immunostaining of tail epidermal sheets

Tail skin was cut into 5 mm² pieces and incubated in 20 mM EDTA at 37°C for 2 hours. The epidermis was separated from the dermis and fixed overnight in 4% PFA at 4°C. Epidermal sheets were blocked for 3 hours in PBS containing 1% BSA, 2.5% NDS, 2.5% NGS, and 0.8% Triton. Then, the samples were incubated with primary antibodies against K10 (Abcam, 1:500) and K36 (Proteintech, 1:300) overnight at RT, followed by Alexa Fluor-conjugated secondary antibodies (Invitrogen, 1:500). The M.O.M. kit (Vector Laboratories) was used to reduce background from mouse-derived antibodies. Nuclei were counterstained with Hoechst 33342 (Sigma-Aldrich, 1:1000). Z-stack images were acquired using Nikon A1 and Zeiss LSM 900 confocal microscopes and are presented as maximum-intensity projections. For overview imaging of epidermal sheets, tile-scanning was performed using a Zeiss LSM 980 confocal microscope.

### Quantification and statistical analysis

Epidermal thickness was measured in H&E-stained tail sections from six interscale/scale epidermal structures per mouse, as previously described (Changarathil et al., 2019). K10/K36-positive areas, tdTomato-positive clone numbers, and clonal areas were manually quantified from 4–8 images per mouse (corresponding to 16–32 interscale or scale regions) using projected Z-stack images analyzed in ImageJ (NIH).

Data distribution was assessed using the Shapiro–Wilk test. For comparisons between two groups, an unpaired Student’s *t* test or Mann-Whitney U test was applied for normally distributed or non-normally distributed data, respectively. For multi-group comparisons, a one-way ANOVA or a Kruskal–Wallis test was used as appropriate. Significance was defined as *p* < 0.05.

## Results

### *Il1r2* is preferentially expressed in the slow-cycling epidermal basal population of tail skin

To characterize the transcriptional heterogeneity of epidermal basal cell populations in mouse tail skin, we reanalyzed a published scRNA-seq dataset (GSE205746)(Ghuwalewala et al., 2022). In K5-tTA/pTRE-H2B-GFP mice, doxycycline suppresses H2B-GFP expression, leading to GFP dilution with cell division and enabling the identification of slow-cycling cells as LRCs (Fig. 1A, B) (Tumbar et al., 2004). In the original study, LRCs, mid-LRCs, and non-LRCs in α6-integrin^+^/Sca1^+^ epidermal basal fraction were isolated based on H2B-GFP label retention following a 3-week doxycycline chase.

Initial clustering recapitulated previously described epithelial populations, annotated based on their marker gene expression: *Krt14*, *Krt15*, and *Col17a1* (epidermal basal cells); *Krt10*, *Krt1*, and *Sbsn* (differentiating cells); *Mki67*, *Top2a*, *Cdc20*, and *Mcm6* (proliferating cells); and *Sox9*, *Krt17*, and *Krt79* (infundibulum) (Fig. S1). Proliferating and infundibulum cell clusters were excluded to focus on non-proliferating epidermal populations in the interfollicular epidermis and hair follicle. The remaining cells were sorted into six transcriptionally distinct clusters: interscale basal, scale basal, hair follicle basal, interscale differentiating, scale differentiating, and hair follicle differentiating cells (Fig. 1C). Cluster identities were supported by the distribution of LRC, mid-LRC, and non-LRC fractions. LRCs were enriched in interscale basal and hair follicle basal clusters, whereas non-LRCs and mid-LRCs were enriched in scale basal and scale differentiating clusters (Fig. 1D).

Marker gene analysis further validated cluster identities (Fig. 1E). Both interscale and scale basal clusters expressed canonical basal markers, including *Krt14* and *Col17a1*. Differentiating clusters were distinguished by lineage-specific markers, with interscale cells expressing *Krt1*/*Krt10* and scale cells expressing *Krt31*/*Krt36*. The hair follicle basal cluster was distinguished by *Krt15, Sox9*, and *Lrig1* expression, whereas the hair follicle differentiation cluster was characterized by *Gata6* and *Krt79* expression, consistent with the cells’ lineage identity.

We next examined the expression patterns of established epidermal stem/progenitor markers alongside *Il1r2* across the six clusters (Table 1) (Fig. 1F, 1G). Previously reported markers showed broad or overlapping expression across multiple clusters; however, *Il1r2* showed a selective enrichment in the interscale basal cluster. We performed immunofluorescence staining of tail skin sections to validate this expression pattern at the protein level. Il1r2^+^ cells were specifically localized to the interscale basal layer, beneath the K10^+^ suprabasal layers, and co-expressed the basal marker K5 (Fig. 1H–K). Together, these results establish Il1r2 as a selective marker of a transcriptionally and spatially distinct slow-cycling basal cell population in the interscale epidermis.

### Generation of Il1r2-CreERT2 knock-in mice using the CRIS-PITCh method

Given the selective enrichment of *Il1r2* in the slow-cycling interscale basal population, we generated an Il1r2-CreERT2 knock-in mouse line. Using a CRISPR-Cas9-mediated CRIS-PITCh method, a CreERT2 cassette was inserted at the translational start site of the *Il1r2* locus (Fig. 2A), thereby disrupting the coding sequence while placing CreERT2 expression under control of the *Il1r2* promoter (Fig. 2A).

**Figure 2.**
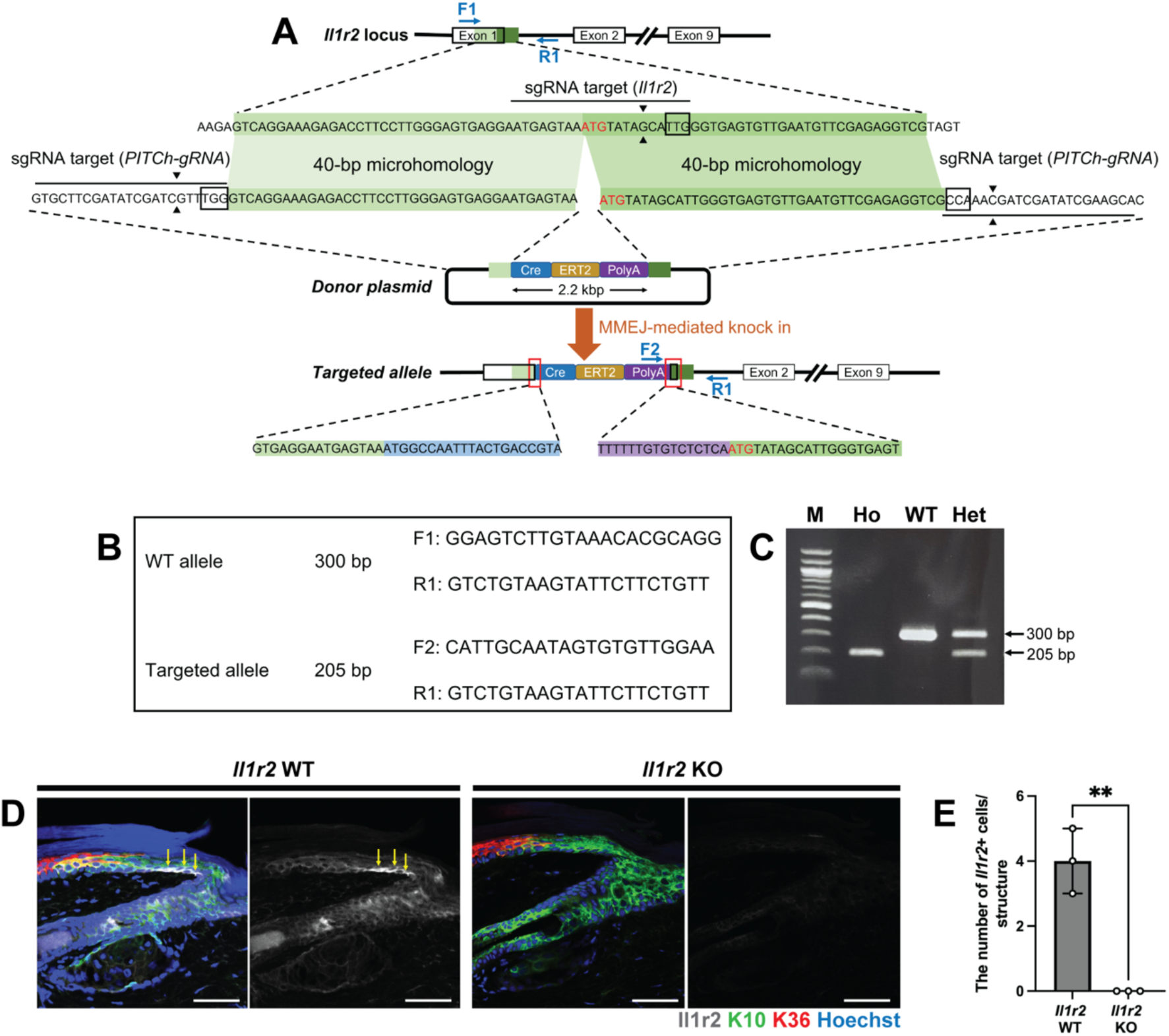
Generation and validation of Il1r2-CreERT2 knock-in mice. **(A)** Schematic of the knock-in strategy showing the sgRNA target sequence and insertion site of the CreERT2-polyA cassette at the *Il1r2* locus. Blue arrows indicate the primer binding sites for genotyping. **(B)** Primer sequences used for genotyping PCR of Il1r2-CreERT2 knock-in mice. The 300-bp product corresponds to the wild-type allele, and the 205-bp product corresponds to the targeted allele. **(C)** Representative PCR results identifying wild-type (WT), heterozygous (Het), and homozygous (Ho) Il1r2-CreERT2 knock-in mice. M, 100-bp DNA ladder. **(D)** Immunostaining of tail skin sections for Il1r2, K10 (interscale marker), and K36 (scale marker) in wild-type and *Il1r2* knockout (KO) mice. **(E)** Quantification of the number of Il1r2^+^ cells per epidermal structure. Yellow arrows indicate Il1r2^+^ cells. Scale bars: 50 µm. Data are presented as mean ± SD. Each dot represents one mouse. Significance was assessed using an unpaired Student’s *t* test. ∗∗*p* < 0.01.

Correct insertion was confirmed by allele-specific PCR: wild-type and knock-in alleles yielded 300-bp and 205-bp PCR amplicons, respectively (Fig. 2B, C). Of the 109 founder mice screened, 26 (23.85%) were identified as Cre-positive by PCR genotyping, indicating efficient CRIS-PITCh targeting. Consistent with the disruption of the *Il1r2* coding sequence, homozygous mice lacked Il1r2 protein expression in the interscale basal layer, as confirmed by immunofluorescence staining (Fig. 2D, E). These results demonstrate the successful generation of the Il1r2-CreERT2 knock-in mouse line that enables tamoxifen-inducible lineage tracing of Il1r2-expressing cells and provides a functional *Il1r2* knockout model in homozygous mice.

### *Il1r2* is dispensable for epidermal homeostasis under steady-state conditions

We next assessed whether *Il1r2* deletion affects overall health, skin structure, and epidermal stem cell organization. *Il1r2* knockout (KO) mice were viable and showed no overt abnormalities in appearance compared with wild-type (WT) controls (Fig. 3A). Body weight was comparable between *Il1r2* KO and WT mice in both sexes (Fig. 3B), indicating that *Il1r2* deletion does not impair general health.

**Figure 3.**
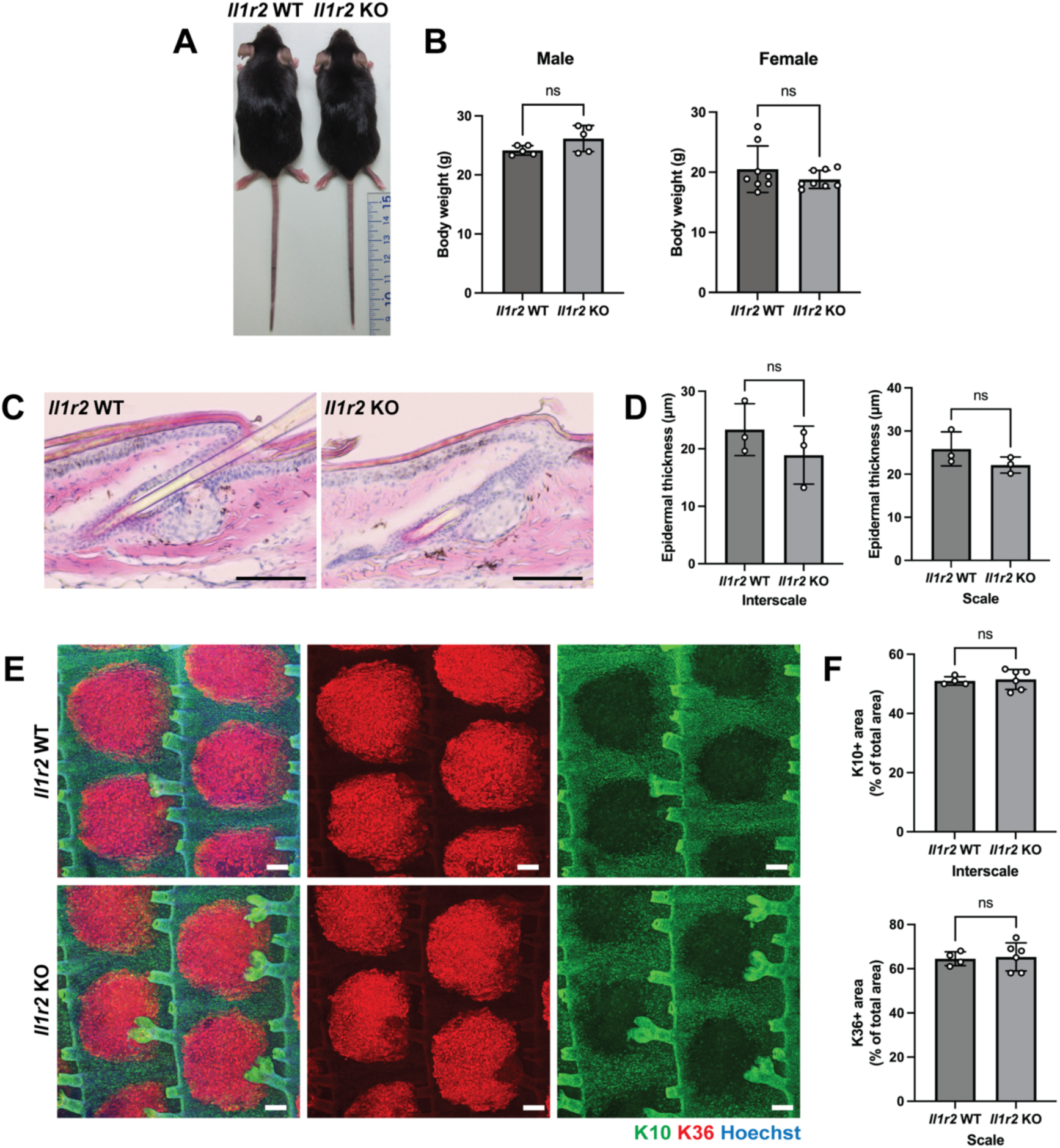
Epidermal structure and stem cell organization are preserved in *Il1r2* knockout mice. **(A)** Representative photographs of *Il1r2* wild-type (WT) and knockout (KO) mice at 2–3 months of age. **(B)** Body weight of *Il1r2* WT and KO mice in males and females. **(C, D)** Hematoxylin and eosin staining of sagittal tail skin sections (C), and quantification of epidermal thickness in the interscale and scale regions (D). Scale bars: 100 µm. **(E)** Whole-mount immunostaining of tail epidermal sheets for K10 (interscale marker) and K36 (scale marker). **(F)** Quantification of K10^+^ and K36^+^ areas. Scale bars: 100 µm. Data are presented as the mean ± SD. Each dot represents one mouse. Significance was assessed using an unpaired Student’s *t* test. ns, not significant.

Histological analyses of tail skin sections revealed no significant differences in epidermal thickness, stratification, or overall structure between *Il1r2* KO and WT mice (Fig. 3C, D). Whole-mount immunostaining of epidermal sheets further showed that the K10^+^ interscale and K36^+^ scale regions were preserved with comparable size and spatial organization in *Il1r2* KO mice (Fig. 3E, F). These results indicate that *Il1r2* is dispensable for the formation or maintenance of epidermal structure and stem cell organization during homeostasis.

### Il1r2-CreERT2 preferentially labels the interscale epidermal population

To examine the distribution of Il1r2-CreERT2 labeled cells, we crossed Il1r2-CreERT2 knock-in mice with Rosa-tdTomato reporter mice. In Il1r2-CreERT2/Rosa-tdTomato mice, tamoxifen administration induced Cre-mediated recombination in Il1r2-expressing cells, resulting in tdTomato expression in labeled cells and their progeny (Fig. 4A). Adult mice (2–3 months) received a single intraperitoneal dose of tamoxifen, and tail skin was collected after a 2-week chase to capture the initial labeling pattern (Fig. 4B).

**Figure 4.**
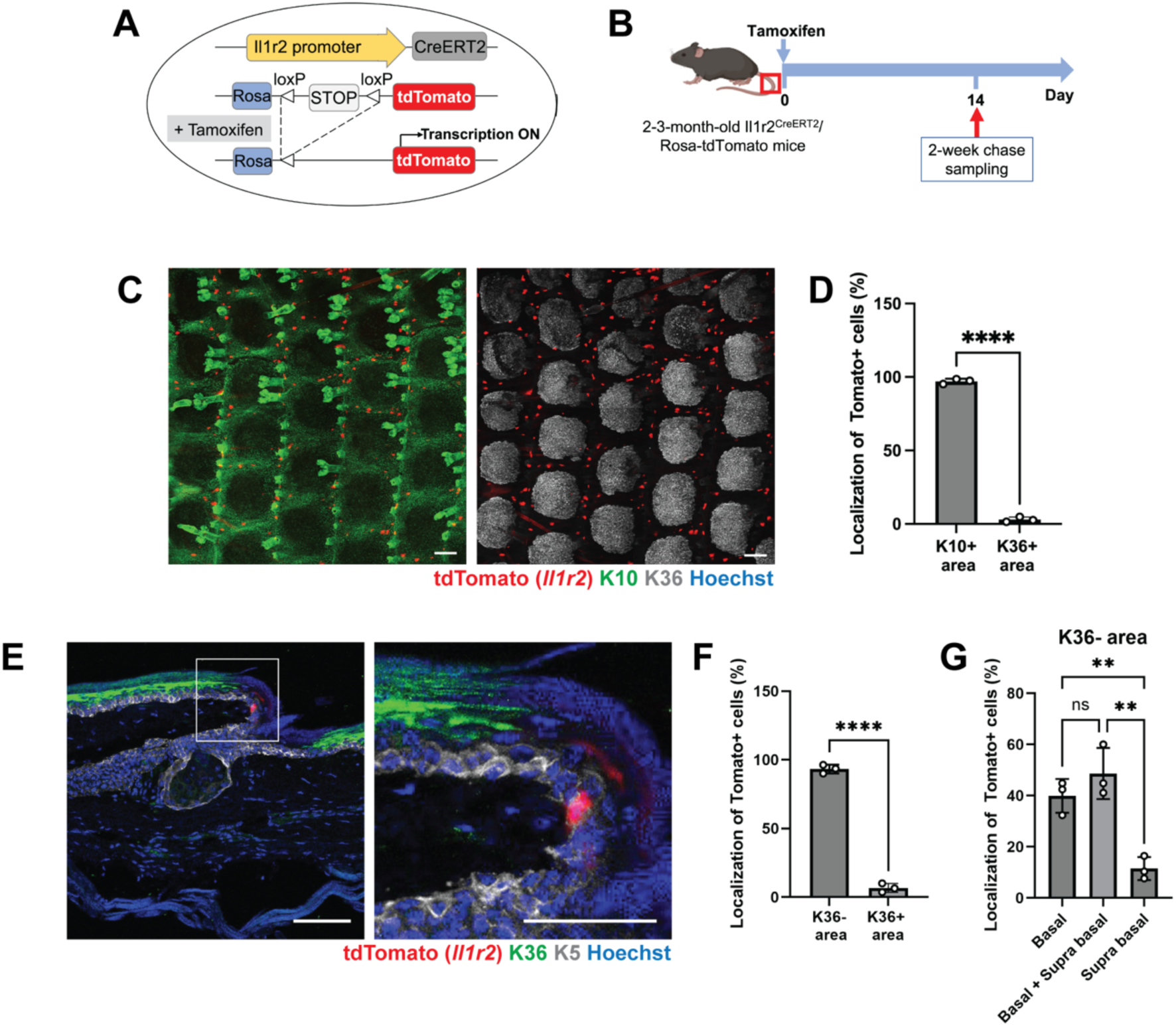
Spatial distribution of Il1r2-CreERT2 labeled cells in tail skin. **(A)** Schematic of the tamoxifen-inducible CreERT2 system used for lineage tracing. **(B)** Experimental timeline. **(C)** Whole-mount immunostaining of tail epidermal sheets showing Il1r2-CreERT2^+^ clones co-stained with K10 (interscale marker) and K36 (scale marker). **(D)** Quantification of the percentage of Il1r2-CreERT2^+^ clones localized within the K10^+^ interscale and K36^+^ scale regions. **(E)** Immunostaining of tail skin sections showing Il1r2-CreERT2^+^ (tdTomato^+^) clones co-stained with K5 (basal layer marker) and K36 (scale marker). **(F)** Quantification of the percentage of Il1r2-CreERT2^+^ clones localized within the K36^−^ interscale and K36^+^ scale regions. **(G)** Quantification of clone composition, categorized as basal layer only, basal plus suprabasal layers, or suprabasal layer only. All data are presented as the mean ± SD. Each dot represents an independent biological replicate. Significance was assessed using an unpaired Student’s *t* test for (D, F) and a one-way ANOVA followed by Dunnett’s post hoc test for (G). ∗∗*p* < 0.01; ∗∗∗∗*p* < 0.0001; ns, not significant. Scale bars: 100 µm (C) and 50 µm (F).

Whole-mount immunostaining of epidermal sheets revealed that tdTomato⁺ cells were highly enriched in the K10^+^ interscale region and were rarely detected in the K36⁺ scale regions or hair follicles (Fig. 4C, D), demonstrating a strong spatial bias of Il1r2-CreERT2 labeling. This distribution was consistent with the localization of Il1r2 protein and scRNA-seq data (Fig. 1F–I).

Immunostaining of tail skin sections further confirmed that tdTomato^+^ cells were confined to the K36^-^ interscale region and were largely absent from the K36^+^ scale region (Fig. 4E, F). TdTomato⁺ cells were not detected in the dermis, hair follicles, or other epidermal appendages. Notably, quantitative analysis revealed that approximately 40% of tdTomato⁺ clones in the interscale were located in the basal layer, whereas ∼50% spanned both basal and suprabasal layers, and ∼10% were present in the suprabasal layer (Fig. 4G). In contrast, Il1r2 protein expression was exclusively observed in basal cells (Fig. 1J, K), indicating that Il1r2-expressing cells give rise to differentiating progeny within this 2-week chase period. Together, these results demonstrate that Il1r2-CreERT2 selectively labels basal cells in the interscale region of tail skin, consistent with the known distribution of slow-cycling epidermal basal cells.

## Discussion

The spatial organization of epidermal stem/progenitor cells is thought to enable distinct cell populations to respond differentially to physiological and pathological conditions, including tissue injury, inflammation, and environmental stress (Dumrongphuttidecha et al., 2026; Fan et al., 2026; Ghuwalewala et al., 2022; Phung et al., 2026; Raja et al., 2022; Rognoni & Watt, 2018; Sada et al., 2016; Sánchez-Danés et al., 2016). However, the lack of specific markers has limited the *in vivo* study of these distinct basal populations. In this study, we identify *Il1r2* as a marker enriched in the interscale basal cell population of tail skin. Notably, the Il1r2-CreERT2 model exhibited selective labeling of the interscale basal layer with minimal activity detected in the scale, hair follicles, or dermis, distinguishing it from previously reported CreER lines (Table 1). This finding highlights the utility of this model as a genetic tool for inducible lineage tracing and targeted gene manipulation of the interscale basal cell population. However, whether *Il1r2*-expressing cells possess epidermal stem cell potential remains to be determined. Future studies using long-term lineage tracing will be required to assess the self-renewal capacity and contribution of this population to epidermal maintenance over time.

We generated an Il1r2-CreERT2 knock-in mouse line using a CRIS-PITCh via zygote microinjection (Nakamae et al., 2017; Tanimoto et al., 2022). This approach enables the efficient insertion of relatively large DNA cassettes (∼2.2 kb) into a defined genomic locus without the long homology arms required by conventional homologous recombination in embryonic stem cells. Electroporation-based genome editing is widely used to introduce smaller DNA fragments; however, the insertion of larger gene cassettes typically requires delivery of double-stranded DNA donors via zygote microinjection. Consistent with previous reports of gene cassette knock-in using this method (median efficiency ∼20%)(Tanimoto et al., 2022), we achieved a targeting efficiency of 23.85%, indicating the robustness of this strategy for generating CreERT2 knock-in alleles.

Although we followed a similar data preprocessing pipeline to the original study (Ghuwalewala et al., 2022), our cluster annotation strategy differed in two key aspects. First, the original analysis assigned cluster identities using gene signatures from back skin to distinguish LRC and non-LRC populations, which may not fully reflect the transcriptional features or cellular composition of the tail epidermis. Second, hair follicle lineage cells were not clearly separated from interfollicular epidermal populations, potentially introducing unwanted transcriptional heterogeneity within clusters. In our re-analysis, cluster identities were validated by the distribution of experimentally defined LRC, mid-LRC, and non-LRC fractions in combination with marker gene expression. This approach enabled the clearer separation of hair follicle and interfollicular epidermal populations and refined annotation of interscale and scale clusters, leading to the identification of *Il1r2* as a selectively enriched marker for the interscale basal cell population.

IL-1 signaling plays a critical role in inflammatory responses to tissue damage and stress. IL-1α and IL-1β signal through their receptor IL-1R1, whereas IL-1R2 acts as a decoy receptor that limits excessive signaling by sequestering IL-1 ligands (Garlanda et al., 2025). Although *Il1r2* is expressed in the slow-cycling basal population, *Il1r2* KO mice display no overt phenotype, suggesting that IL-1R2 is dispensable for homeostatic tissue maintenance. Instead, its functional importance is likely context-dependent. Dysregulated IL-1 signaling contributes to inflammatory- or age-related pathologies and stem cell impairment (Ishikawa et al., 2025; Morinaga et al., 2021; Phung et al., 2026; Vu et al., 2022). IL-1R2 has been proposed as a potential druggable target due to its ability to attenuate cutaneous inflammation (Calabrese et al., 2022; Rauschmayr et al., 1997). Our Il1r2-CreERT2 model provides a platform to dissect the role of IL-1R2 in defined epidermal subpopulations, enabling simultaneous lineage tracing and conditional loss-of-function analysis in contexts of tissue remodeling, stress, and disease.

## Acknowledgments

We thank the Center for Animal Resources and Development (CARD), Kumamoto University, and the Laboratory of Embryonic and Genetic Engineering at Kyushu University for support with animal facilities. We also thank the International Core-facility of Advanced Life Science at Kumamoto University and the Research Promotion Unit at Kyushu University for core facility support. We thank Dr. H. Miura (Tokai University), T. Keida, T. Kosasih (Kumamoto University), and N. Imai (Kyushu University) for technical assistance with mouse experiments. We also thank H. Takizawa and T. Takeo (Kumamoto University) for valuable discussions and critical feedback.

This work was supported by the Project for Regenerative, Cellular Medicine and Gene Therapies (JP26bm1123052) and AMED-PRIME (JP21gm6110016) from the Japan Agency for Medical Research and Development (AMED) (to A.S.); and by Grant-in-Aid for Scientific Research (B) (JP20H03266, JP24K02035) (to A.S.), Grant-in-Aid for Early-Career Scientists (JP22K15126) (to N.T.K.N.), Grant-in-Aid for Challenging Research (Exploratory) (JP24K21973) (to A.S.) from the Japan Society for the Promotion of Science (JSPS). This work was also supported by The Naito Foundation, The Takeda Science Foundation, The Sumitomo Foundation, The Mochida Memorial Foundation for Medical and Pharmaceutical Research, Lydia O’Leary Memorial Pias Dermatological Foundation, and Koyanagi Foundation (all to A.S.). H.M.P. was supported by a scholarship from the Ministry of Education, Culture, Sports, Science, and Technology (MEXT). I.N. was supported by the SPRING program from the Japan Science and Technology Agency (JST) (JPMJSP2136). This work was supported in part by the MEXT Cooperative Research Project Program, Medical Research Center Initiative for High Depth Omics, and CURE: JPMXP1323015486 for MIB, Kyushu University. The infrastructure of Omics Science Center Secure Information Analysis System (OASIS; https://sis.bioreg.kyushu-u.ac.jp/), Medical Institute of Bioregulation at Kyushu University, provides a part of the computational resource.

## Author contributions

A.S. conceived and designed the study. N.T. and K.A. performed zygote microinjection to generate the Il1r2-CreERT2 knock-in mouse line and provided technical advice on mouse design and evaluation. N.T.K.N. and A.S. designed the mouse model and experimental strategy, and N.T.K.N. performed initial screening and validation of the mouse line. H.M.P. performed mouse phenotype analyses and quantification, including *Il1r2* knockout and lineage-tracing experiments. I.N. and T.F. performed the initial scRNA-seq analysis. T.F., Y.O., and G.G. contributed expertise in omics data analysis. H.M.P. performed the reanalysis of scRNA-seq data and contributed to data interpretation and visualization. A.S. provided resources and supervised the project. A.S. and N.T.K.N. acquired funding. H.M.P. wrote the original draft of the manuscript. A.S. reviewed and edited the manuscript. All authors approved the final version of the manuscript.

## Declaration of generative AI and AI-assisted technologies in the writing process

Generative AI (ChatGPT, Gemini, and Claude) was used to support language editing for clarity and readability during manuscript preparation. All content was reviewed and revised by the authors, who take full responsibility for the accuracy and integrity of the final manuscript.

## Ethical statement

All animal procedures were reviewed and approved by the Animal Care and Use Committee of Kumamoto University and the Animal Experiment Committee of Kyushu University. Experiments were conducted in accordance with institutional guidelines and regulations.

---

**Table 1. Comparison of CreER mouse lines for labeling epidermal stem and progenitor cell populations in mouse tail skin**

**Table 2. Primers used for CreERT2 cassette amplification**

**Table 3. gRNA sequences used for *Il1r2* targeting**

## Supplemental information

**Figure S1.**
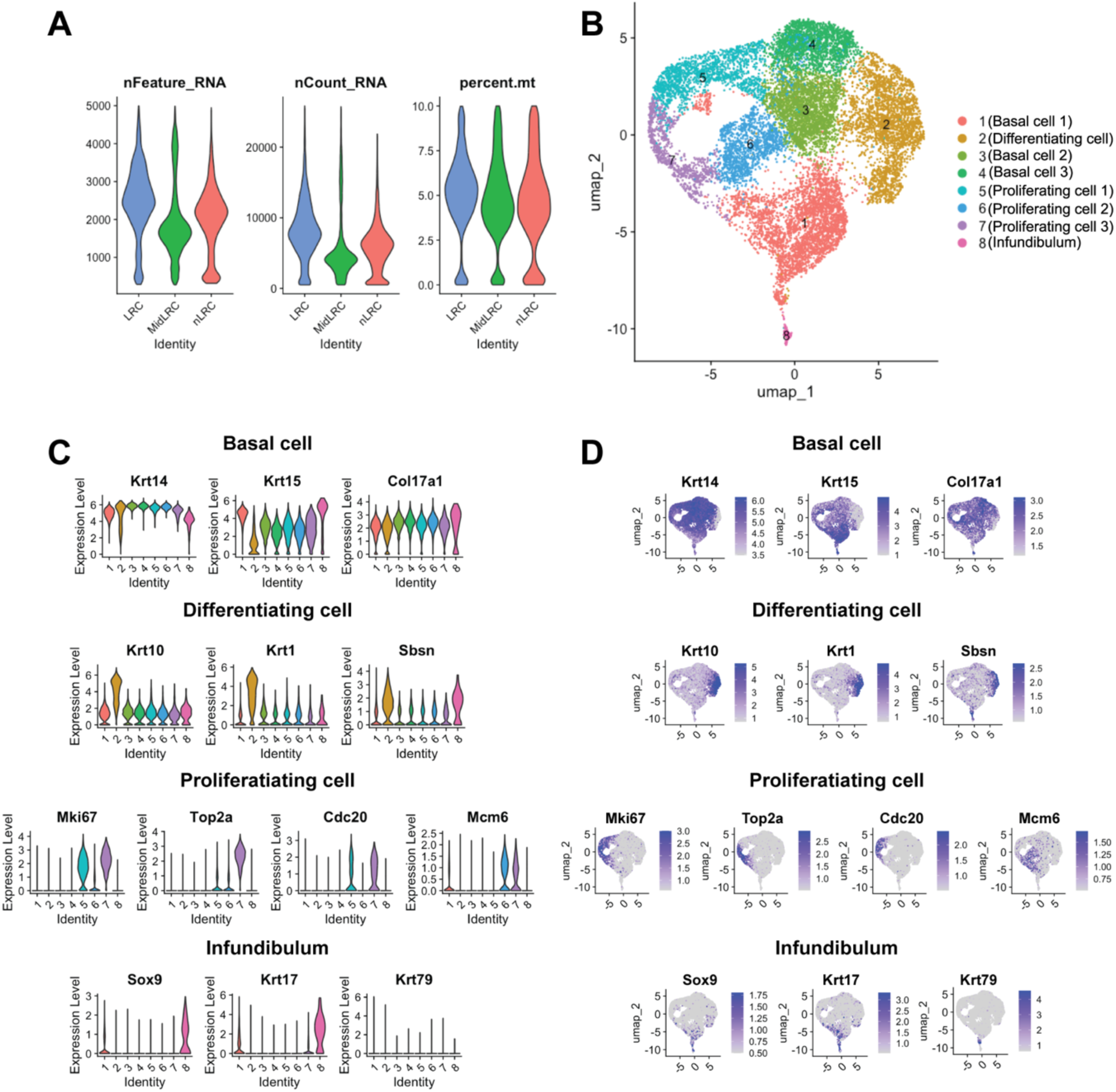
scRNA-seq analysis including infundibulum and proliferating cells prior to reclustering. **(A)** Violin plots showing quality control metrics, including mRNA count, gene count, and mitochondrial gene percentage, for tail skin sorted cells from two biological replicates used in scRNA-seq analysis. **(B)** UMAP plot showing cell clusters identified based on established marker gene expression. **(C)** Violin plots showing normalized expression values of marker genes used to assign cell identities. **(D)** Feature plots showing normalized expression values of marker genes across clusters.

